# A genetic screen to identify deubiquitinases as regulators of IRF7

**DOI:** 10.1101/2025.09.09.675186

**Authors:** Shumin Fan, Pracheta Sengupta, Karan Chawla, Manoj Veleeparambil, Ritu Chakravarti, Saurabh Chattopadhyay

**Author notes:** These authors contributed equally to the work (Shumin Fan, Pracheta Sengupta).

## Abstract

Virus infection rapidly induces the production and secretion of interferons (IFNs), amplifying antiviral responses in infected and neighboring uninfected cells. IFN regulatory factor 7 (IRF7), the ‘master transcription factor,’ is pivotal in IFN induction, particularly in myeloid cells. Ubiquitination of IRF7 is essential for its transcriptional activation; however, the underlying molecular mechanisms remain poorly understood. We hypothesized that deubiquitinases (DUBs) act as endogenous regulators of IRF7 activity and conducted a genetic screen using a human DUB-targeted siRNA library. This screen identified USP2 as a positive regulator and OTUD5 as a negative regulator of IRF7 activity. OTUD5, an inducible DUB, physically interacted with IRF7 and inhibited its K63-linked ubiquitination, thereby suppressing IRF7 activation. Conversely, USP2 promoted IRF7 activity by binding to IRF7 and removing K27-linked ubiquitin chains, which we found to be inhibitory. Specifically, K27-linked ubiquitination impeded phosphorylation of IRF7, a critical step for its activation. Collectively, our genetic screen and mechanistic studies uncovered USP2 and OTUD5 as novel modulators of IRF7 function, providing new insights into the regulation of antiviral immunity.

## Introduction

The type I interferon (IFN) system represents a crucial first line of defense against viral infections and inflammatory insults. Type I IFNs, a class of cytokines primarily secreted in response to viral infection or tissue injury, play a central role in initiating and shaping both innate and adaptive immune responses (1, 2). Their production is triggered by the recognition of pathogen-associated molecular patterns (PAMPs) and damage-associated molecular patterns (DAMPs) by pattern recognition receptors (PRRs). Upon activation, PRRs initiate intracellular signaling cascades that converge on interferon regulatory factors (IRFs), a family of transcription factors critical for the induction of IFNs and other inflammatory mediators (3–5). Among the nine IRF family members identified in mammals, five—IRF1, IRF3, IRF5, IRF7, and IRF8— function as positive regulators of PRR-induced type I IFN responses(6). IRF3 and IRF7, in particular, are structurally related and serve as key transcriptional regulators of type I IFN production downstream of cytosolic RNA sensors (RIG-I-like receptors), DNA sensors (cGAS), and endosomal Toll-like receptors (TLRs). IRF5, while also capable of inducing type I IFNs, plays a prominent role in driving the expression of proinflammatory cytokines such as IL-6, TNF-α, and IL-12. Thus, the interplay among IRF3, IRF5, and IRF7 is essential for determining the magnitude and specificity of the innate immune response (5, 7).

Dysregulation of IRFs is associated with a wide range of pathological conditions, including autoimmune disorders, cancer, and metabolic diseases. In autoimmune diseases such as systemic lupus erythematosus (SLE) and rheumatoid arthritis (RA), aberrant IRF activity leads to excessive IFN production, contributing to chronic inflammation and tissue damage. Conversely, in cancer, IRFs may either support anti-tumor immunity or facilitate immune evasion and tumor progression, depending on the context. For instance, IRF1 and IRF7 have been shown to suppress tumor metastasis. In metabolic disorders, IRF-mediated inflammation may exacerbate insulin resistance and metabolic dysfunction (5, 8). These findings highlight the importance of tightly regulated IRF activity in maintaining immune homeostasis and underscore the potential of IRFs as therapeutic targets.

IRF7 is considered the “master regulator” of type I IFN responses, particularly in immune cells, and plays a pivotal role in the amplification of IFN signaling. While IRF7 expression is inducible by IFNs and PRR signaling, its transcriptional activity depends on posttranslational modifications, primarily phosphorylation. Upon activation by PRRs, adaptor proteins such as MyD88, TRIF, MAVS, and STING recruit the kinases IKKε and TBK1, which phosphorylate IRF7 at key serine residues (e.g., Ser477 and Ser479). This phosphorylation facilitates IRF7 dimerization and nuclear translocation, enabling the transcription of IFN-α and other ISGs (9). In addition to phosphorylation, ubiquitination serves as a critical regulatory mechanism of IRF7 activity. Ubiquitin (Ub), a 76-amino acid protein, can form polyubiquitin chains through any of its seven lysine residues (K6, K11, K27, K29, K33, K48, and K63), with distinct chain types imparting different functional outcomes on target proteins (10, 11). For example, K48-linked ubiquitination commonly targets proteins for proteasomal degradation, whereas K63-linked ubiquitination can enhance protein activation and signaling. In Epstein-Barr virus (EBV) infection, the viral protein LMP1 promotes IRF7 activation via K63-linked ubiquitination at Lys444, Lys446, and Lys452, mediated by the E3 ligase TRAF6. This modification facilitates the recruitment of IKKε, linking IRF7 ubiquitination to its phosphorylation and activation (12–16). However, excessive accumulation of K63-linked ubiquitinated IRF7 (Ub^63^-IRF7) can lead to sustained IFN production, contributing to immunopathology and autoimmunity. Therefore, deubiquitination plays a vital role in modulating IRF7 activity and maintaining immune balance. Deubiquitinases (DUBs) are a class of enzymes that remove ubiquitin chains from proteins, thereby regulating protein turnover, signaling activity, and functional outcomes. Depending on their specificity, DUBs can either enhance or suppress the activity of their targets by selectively cleaving specific ubiquitin linkages (17, 18). In the context of EBV infection, LMP1 recruits the DUB A20 to remove K63-linked ubiquitin chains from IRF7, thereby dampening IRF7 activation and IFN responses (19). DUBs are also essential for maintaining the ubiquitin-proteasome system (UPS) and ensuring proper recycling of free ubiquitin (20). They are broadly classified into four families: ubiquitin-specific proteases (USPs), ubiquitin C-terminal hydrolases (UCHs), ovarian tumor proteases (OTUs), and JAMM metalloproteases (21).

Despite their importance, the specific roles of DUBs in regulating mammalian IRF7 remain poorly understood. In this study, we sought to identify DUBs that modulate IRF7 activity by performing an siRNA-based screen targeting human DUBs. Our screen revealed two novel IRF7 regulators: USP2, which acts as an activator, and OTUD5, which functions as an inhibitor. Mechanistically, USP2 enhances IRF7 activity by removing inhibitory K27-linked ubiquitin chains, whereas OTUD5 attenuates IRF7 function by cleaving activating K63-linked chains. These findings provide new insights into the posttranslational regulation of IRF7 and offer potential therapeutic targets for modulating IFN responses in viral and inflammatory diseases.

## Materials and Methods

### Antibodies

The following antibodies were used: anti-actin (Sigma-Aldrich #A5441), anti-FLAG [Cell Signaling Technology (CST) #2368 and Sigma-Aldrich #F1804], anti-HA (CST #3724S), anti-HDAC1 (CST #34589), anti-IRF3 and anti-IFIT [as previously described (22–24)], anti-IRF7 (CST #39659 and Santa Cruz Biotechnology #sc-74471), anti-phospho-IRF7 (CST #24129), anti-OTUD5 (Invitrogen #PA5-20611), anti-TRAF6 (Santa Cruz Biotechnology #sc-7221), anti-USP2 (Proteintech #10392-1-AP and Invitrogen #MA5-37724), anti-ubiquitin (CST #58395S), and anti-V5 (CST #13202).

### Cell Lines, Plasmids, and Viruses

HEK293T/17 (CRL-11268) and RAW264.7 (TIB-71) cells were obtained from ATCC; RAW-Lucia ISG (RAWL-ISG) cells were from Invivogen. Cells were cultured in DMEM supplemented with 10% fetal bovine serum and 1% penicillin/streptomycin at 37°C in 5% CO . Mycoplasma removal agent (MP Biomedicals) was routinely used. Plasmids included HA-Ub-K27O (Addgene #22903), HA-Ub-K63O (Addgene #17606), OTUD5 (Sino Biologicals #HG22452-CF), and USP2 (Sino Biologicals #HG12713-UT). Sendai virus (SeV Cantell) was from Charles River Laboratories. GFP-tagged VSV and IAV strain PR8 were previously described (24, 25).

### Gene Knockout and Overexpression Cell Lines

IRF3 knockout (KO) HEK293T cells (HEK-KO) were generated using CRISPR/Cas9 as described (23). V5-tagged IRF7 was used to generate HEK-KO.IRF7 stable cell lines. Knockout and overexpression were confirmed by immunoblotting.

### Cloning and Mutagenesis

Mouse IRF7 (Origene: NM_016850) was subcloned into the pLVX-IRES-puromycin vector (Clontech) to generate V5-tagged wild-type and deletion mutants(23). Constructs were generated using standard molecular biology techniques and verified by sequencing. V5-tagged lysine-to-arginine (KR) point mutants of IRF7 were synthesized by GenScript.

### Cell Lysis and Immunoblotting

Cells were lysed in 50 mM Tris-HCl (pH 7.4), 150 mM NaCl, 0.1% Triton X-100, 1 mM sodium orthovanadate, 10 mM NaF, 10 mM β-glycerophosphate, 5 mM sodium pyrophosphate, and protease/phosphatase inhibitors. Lysates were sonicated and centrifuged (30 min), and protein concentrations were measured using Bradford reagent (Bio-Rad #500–0006). Equal amounts of protein were analyzed via SDS–PAGE and immunoblotting.

### Co-Immunoprecipitation and Ubiquitination Assays

For co-immunoprecipitation (co-IP), cells were lysed in EPPS buffer with protease/phosphatase inhibitors (Roche), subjected to freeze-thaw cycles, and centrifuged. For ubiquitination assays, cells were lysed in Tris buffer (50 mM, pH 7.4) with 150 mM NaCl, 0.2% Triton X-100, 0.5% SDS, 1 mM sodium orthovanadate, 10 mM NaF, 10 mM β-glycerophosphate, 5 mM sodium pyrophosphate, protease/phosphatase inhibitors (Roche), and MG132. Lysates were subjected to freeze thaw cycles and centrifuged at 13,000 rpm. Immunoprecipitation was performed using V5-agarose beads (Sigma #SAE0203) overnight, followed by sequential washing with EPPS and RIPA buffers. Bound proteins were eluted by boiling in SDS buffer with 2-mercaptoethanol and analyzed by SDS–PAGE and immunoblotting. Nuclear fractions were isolated as previously described (23, 24).

### Cellular Signaling and Viral Infections

TLR3 signaling was induced by adding poly I:C (25 µg/mL) to culture medium for 8 h. RLR signaling was activated by transfecting cells with poly I:C using Lipofectamine 2000 (Thermo Fisher Scientific), denoted as pIC+LF, for 8 h unless otherwise noted. TLR9 signaling was activated by adding CpG ODN (10 µg/ml) to the culture media (26, 27), and TLR7 signaling was stimulated by adding R848 (10 µg/ml) to the culture media (28). For virus infections, cells were adsorbed with SeV (MOI 10), IAV (MOI 5), or GFP.VSV (MOI 0.1) in serum-free DMEM for 2 h, then washed and incubated in complete medium. Cells were harvested at the indicated time points for analysis by qRT-PCR or immunoblotting.

### Proximity Ligation Assay and Confocal Microscopy

Cells grown on coverslips were fixed with 4% paraformaldehyde (Electron Microscopy Sciences #15710) and permeabilized with 0.2% Triton X-100 (Thermo Fisher Scientific #9002–93-1). Cells were immunostained with anti-IRF7 and anti-OTUD5 or anti-USP2 antibodies, followed by Duolink proximity ligation assay (Sigma-Aldrich: DUO92008-3, DUO92004, DUO92002) per the manufacturer’s instructions. Coverslips were mounted using VectaShield with DAPI (Vector Laboratories #H-1200) and imaged using a Zeiss microscope (Zen Blue) and Zeiss Zen 3.7 software.

### siRNA Transfection

siRNAs targeting human TRAF6, OTUD5, USP2, and a SMARTpool siRNA library for human deubiquitinating enzymes (10 μM, Horizon Discovery #G-104705, Lot 19124) were transfected using DharmaFECT-1 reagent (Horizon Discovery). A non-targeting siRNA (Thermo Scientific #D-001810-10-05) was used as a control. After 48 h, cells were used for co-IP, ubiquitination, or qRT-PCR analyses, as described in the figure legends.

### Primary Cell Culture

Bone marrow-derived macrophages (BMDMs) were isolated from wild-type male and female mice using established protocols (26, 29). All animal procedures were approved by the Institutional Animal Care Committees of the University of Toledo (Protocol #108668) and the University of Kentucky (Protocol #2023-4348).

### DUB Screening Protocol for IRF7 Regulation

HEK-KO.IRF7 cells were transfected with control (siNT) or DUB-specific siRNAs (siDUBs) for 60 h. Cells were infected with SeV (MOI 10) for 8 h, and IFIT3 expression was analyzed by immunoblot. Twenty-four candidate DUBs identified in the primary screen were further tested in secondary screening via qRT-PCR for IFNA expression. Fourteen DUBs were selected for tertiary screening, in which Co-IP and qRT-PCR were used to assess IRF7 ubiquitination and IFIT1 induction, respectively. An overview of the screening workflow is presented in Fig. 2A, and results are summarized in Fig. 3F.

**Figure 1.**
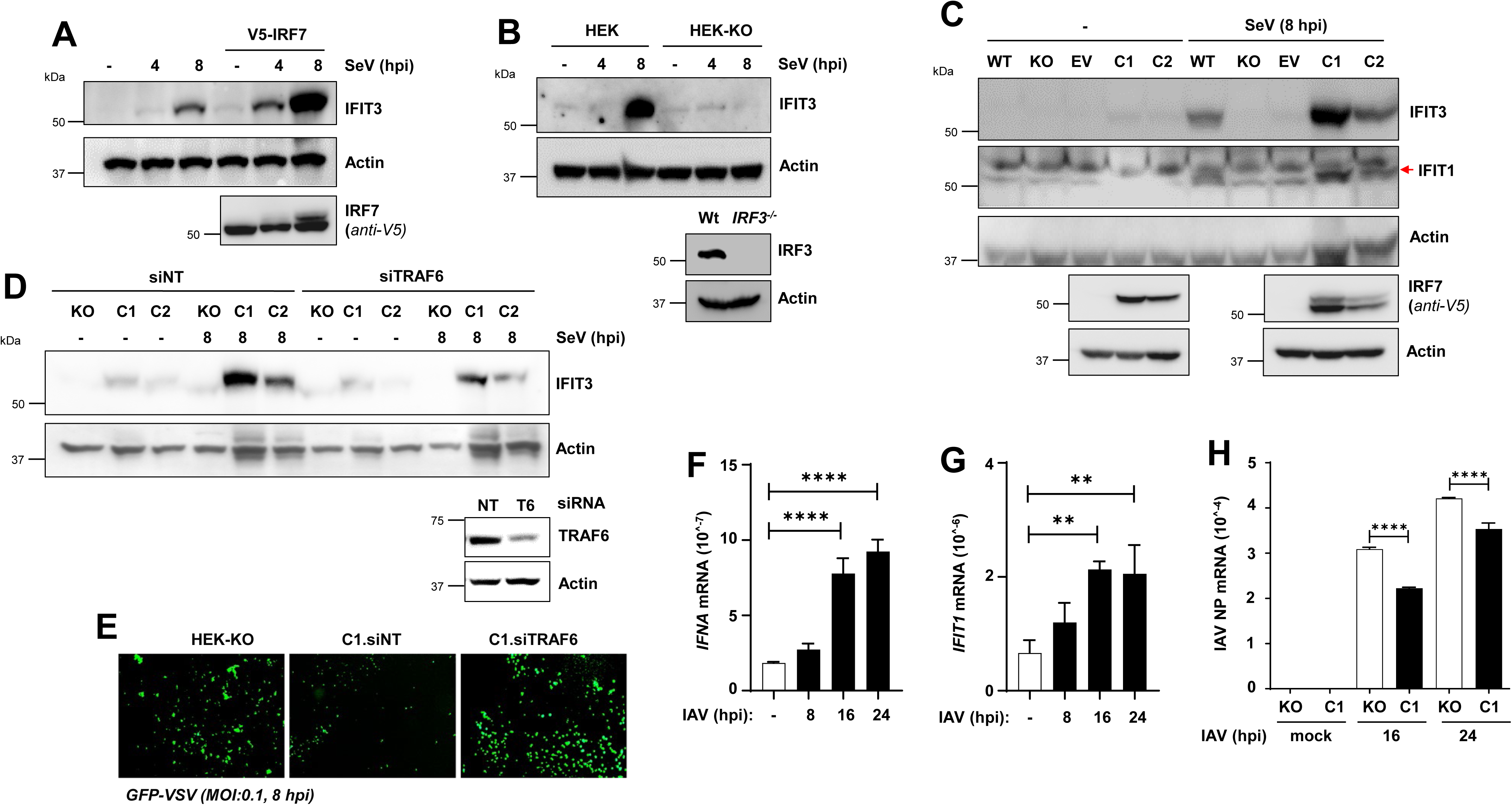
Establishing a functional screening model for IRF7-specific regulators in the absence of IRF3. **(A)** HEK293T (HEK) cells were transfected with V5-tagged IRF7 and infected with SeV for the indicated times. IFIT3 and IRF7 expression were analyzed by immunoblotting. **(B)** Wild-type (WT) and IRF3-deficient (HEK-KO) HEK293T cells (HEK) were infected with SeV as indicated, and IFIT3 induction was assessed by immunoblot. IRF3 protein levels are shown in the lower panel. **(C)** HEK-KO cells were transfected with V5.IRF7, and two stable clones (C1 and C2) were tested for IFIT3 and IFIT1 induction upon SeV infection. IRF7 expression is shown in the lower panels. **(D)** Clones C1 and C2 were transfected with non-targeting (NT) or TRAF6-specific siRNAs and analyzed for IFIT3 induction upon SeV infection by immunoblot. TRAF6 knockdown is shown below. **(E)** HEK-KO and HEK-KO.IRF7 (C1) cells were transfected with NT or TRAF6-specific siRNAs and infected with GFP.VSV. Representative fluorescence microscopy images are shown. **(F–H)** HEK-KO.IRF7 cells were infected with Influenza A Virus (IAV; MOI:10) for the indicated times. *IFNA*, *IFIT1*, and viral NP mRNA levels were quantified by qRT-PCR. EV, empty vector; NT, non-targeting. The data represent mean ± SEM (F-H), ** p<0.001, **** p<0.0001.

**Figure 2.**
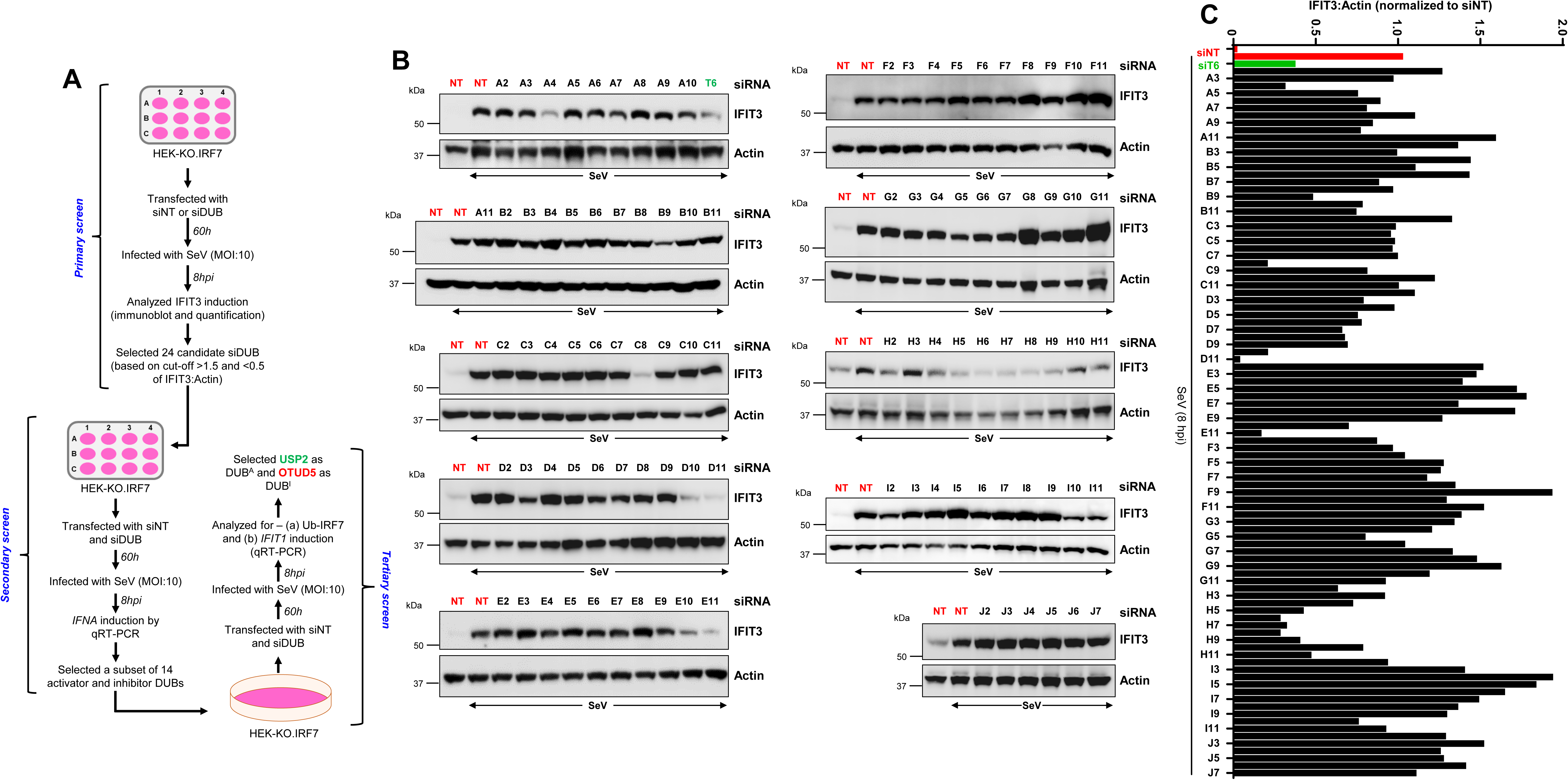
Primary siRNA screen of human deubiquitinases (DUBs) to identify IRF7 regulators. **(A)** Schematic overview of the DUB siRNA screening workflow in HEK-KO.IRF7 cells. Cells were transfected with siRNAs, infected with SeV, and analyzed for IFIT3 induction by immunoblot (primary screen). The shortlisted candidates from the primary screen were followed by secondary and tertiary screens, as indicated. **(B)** HEK-KO.IRF7 cells were transfected with individual DUB siRNAs (10 µM) for 60 hours, followed by SeV infection (MOI:10) for 8 hours. IFIT3 expression was analyzed by immunoblot. TRAF6 siRNA served as a positive control; NT siRNA was included in each batch. **(C)** Densitometric quantification of IFIT3 induction from (B), normalized to actin using ImageJ. NT, non-targeting.

**Figure 3.**
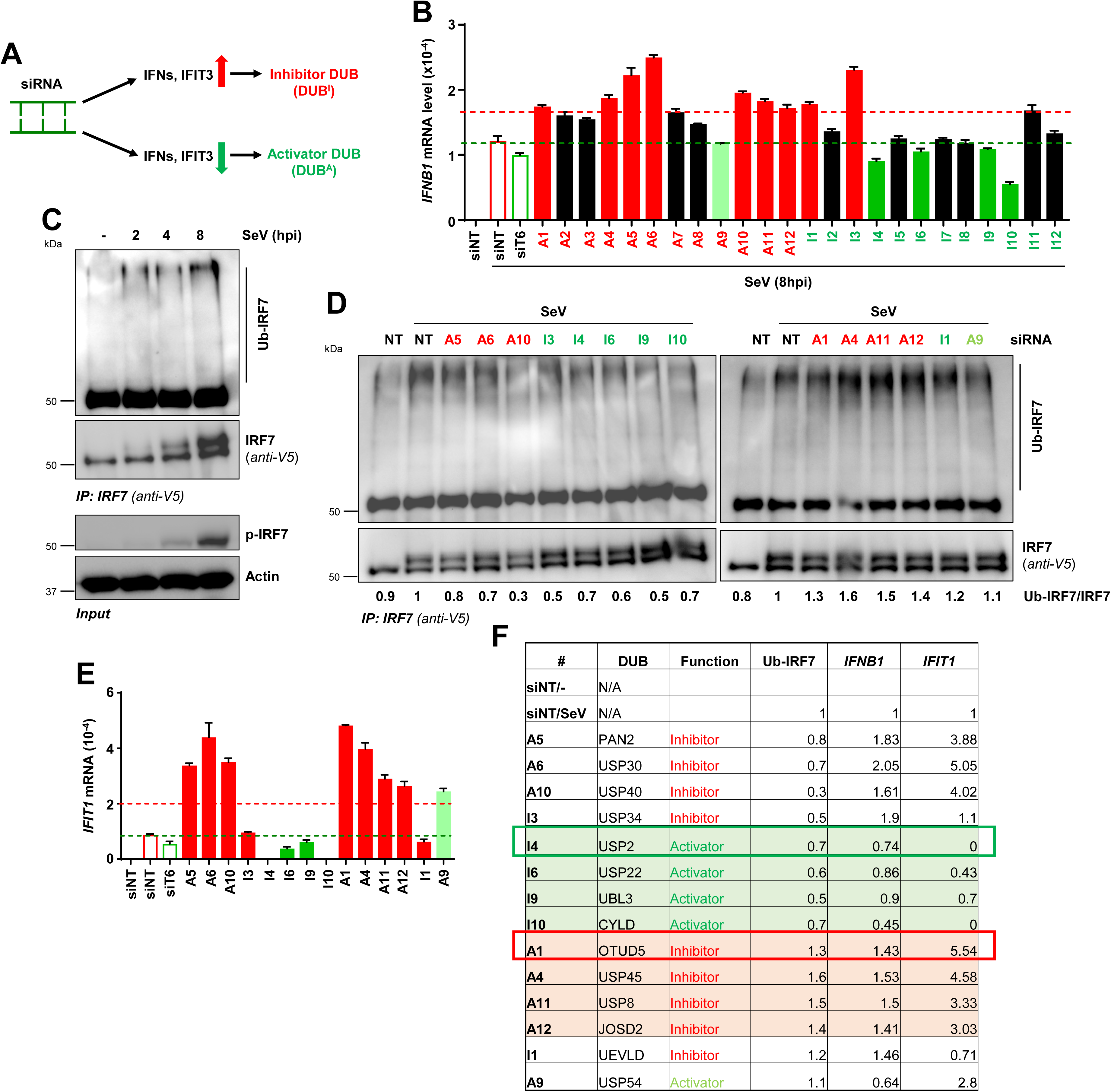
Secondary and tertiary screening identify USP2 and OTUD5 as IRF7-specific regulators. **(A)** Criteria used to select candidate DUBs for follow-up screening. **(B)** In secondary screening, selected DUB siRNAs were transfected into HEK-KO.IRF7 cells and *IFNB1* mRNA levels were measured by qRT-PCR after SeV infection. **(C)** HEK-KO.IRF7 cells were infected with SeV for the indicated times. IRF7 ubiquitination was analyzed by immunoprecipitation using anti-V5 antibody followed by anti-ubiquitin (Ub) immunoblot. Input cell lysates were probed for p-IRF7. **(D)** Candidate DUB siRNAs from (B) were tested for IRF7 ubiquitination using the assay in (C), and relative Ub-IRF7 levels were quantified by Image J. **(E)** *IFIT1* mRNA induction in response to SeV infection was quantified by qRT-PCR in cells transfected with candidate DUB siRNAs. **(F)** Summary of secondary and tertiary screen results. Highlighted rows indicate DUB candidates selected for further study. NT, non-targeting.

### RNA Isolation and qRT-PCR

RNA was extracted using TRIzol (Sigma #T9424) and treated with DNase I (Promega). cDNA synthesis was performed using random hexamers and the ImProm-II Reverse Transcription Kit (Promega). qRT-PCR was carried out using Radiant SYBR Green PCR mix (Alkali Scientific Inc.) on Roche LightCycler 96 and Applied Biosystems 7500 Fast Dx instruments. Data were analyzed using LightCycler 480 v1.5 and SDS v1.4.1 software. Expression levels were normalized to 18S rRNA and plotted using GraphPad Prism 10.

### Quantification and Statistical Analysis

The data represent at least three independent experiments, and statistical analyses were conducted using GraphPad Prism 10, incorporating both biological and technical replicates. Two-tailed unpaired Student’s t-tests were used for two-group comparisons, and one-way ANOVA for multiple-group comparisons. The p-values < 0.05 were considered statistically significant.

## Results

### Establishing a Human Cell Model to Screen Regulators of IRF7 Transcriptional Activity

Early antiviral responses are primarily mediated by IRF3, which is expressed ubiquitously and activated rapidly upon infection. However, many viruses target IRF3 for degradation to evade host immune defenses. To investigate whether IRF7 can functionally compensate for IRF3, we established a HEK293T (HEK) cell model with ectopic IRF7 expression, mimicking its physiological induction during infection. Compared to parental cells, IRF7-expressing cells showed enhanced induction of the antiviral gene *IFIT3* following Sendai virus (SeV) infection (Fig. 1A), indicating that IRF7 is transcriptionally active in this model. To assess IRF7’s ability to replace IRF3, we generated an IRF3 knockout HEK cell line (HEK-KO) using CRISPR/Cas9 (Fig. 1B), which, as expected, failed to induce *IFIT3* upon SeV infection. We then ectopically expressed V5-tagged IRF7 (V5.IRF7) in HEK-KO cells and isolated two clones (C1 and C2) with different IRF7 expression levels. Both clones induced *IFIT3* and *IFIT1* upon SeV infection in the absence of IRF3 (Fig. 1C). To determine whether IRF7 activation depended on canonical antiviral signaling, we used siRNA to knock down TRAF6, a known E3 ubiquitin ligase for IRF7. TRAF6 knockdown reduced SeV-induced *IFIT3* expression in both clones (Fig. 1D), confirming pathway engagement. This impaired response correlated with enhanced replication of vesicular stomatitis virus (VSV), as indicated by increased GFP reporter expression (Fig. 1E). We selected clone C1 (HEK-KO.IRF7) for further analysis using influenza A virus (IAV) infection. Compared to HEK-KO cells, HEK-KO.IRF7 cells upregulated IRF7-target genes (*IFNA*, *IFIT1*) (Fig. 1F, G) and suppressed IAV NP mRNA levels (Fig. 1H), confirming antiviral functionality. Together, these data establish HEK-KO.IRF7 as a functional human cell model in which IRF7 can substitute for IRF3 to drive antiviral gene expression and restrict viral replication. This model provides a valuable platform to screen for regulators of IRF7 transcriptional activity.

### A Genetic Screen to Identify Deubiquitinases Regulating IRF7 Transcriptional Activity

The specific ubiquitin (Ub) linkages conjugated to IRF7, and the deubiquitinases (DUBs) responsible for their removal, remain poorly understood. To identify novel DUBs that regulate IRF7 transcriptional activity, we employed the HEK-KO.IRF7 human cell model to conduct a genetic siRNA screen (Fig. 2A). In the primary screen, HEK-KO.IRF7 cells were transfected in a 96-well format with pooled siRNAs (four per gene) targeting individual human DUBs, alongside a non-targeting control (siNT). Following siRNA transfection, cells were infected with Sendai virus (SeV), and IRF7-mediated induction of the antiviral protein IFIT3 was assessed by immunoblotting (Fig. 2B). IFIT3 expression was normalized to actin, and relative levels were quantified against siNT controls (Fig. 2C). As a positive control, siRNA against TRAF6 (siT6), a known upstream activator of IRF7, effectively suppressed IFIT3 induction, validating the screen. Contrary to our expectations, the primary screen revealed both potential positive and negative DUB regulators of IRF7 activity, based on changes in IFIT3:Actin ratios (Fig. 2C). We reasoned that knockdown of an activating DUB would reduce IFIT3 expression, whereas knockdown of an inhibitory DUB would enhance it (Fig. 3A). Using an arbitrary threshold (fold change >1.5 for activators; <0.5 for inhibitors), we shortlisted twelve candidate activator DUBs and twelve inhibitor DUBs for secondary screening. In the secondary screen, we assessed IRF7-dependent *IFNB1* mRNA expression by qRT-PCR in response to SeV infection. This allowed us to narrow the list to seven candidate activators and seven candidate inhibitors (Fig. 3B). These fourteen DUBs were advanced to a tertiary screen that evaluated IRF7 ubiquitination and induction of a second IRF7 target gene, *IFIT1*. To monitor steady-state ubiquitination of IRF7, we developed a biochemical assay to detect ubiquitinated IRF7 (Ub-IRF7) under native conditions, without protease inhibitors, to reflect physiological turnover. SeV infection robustly induced Ub-IRF7 (Fig. 3C). In parallel, IRF7 phosphorylation was also detected, as evidenced by a mobility shift using anti-V5 antibody and confirmation with phospho-serine-specific antibodies (Fig. 3C, bottom panel). Each of the fourteen DUB candidates was individually knocked down by siRNA, and Ub-IRF7 levels were quantified (Fig. 3D). In parallel, *IFIT1* mRNA induction was measured by qRT-PCR (Fig. 3E). Results from the secondary and tertiary screens were integrated to compare candidate DUBs based on four criteria: (a) regulation of Ub-IRF7 levels, (b) impact on *IFNB1* mRNA expression, (c) effect on *IFIT1* mRNA induction, and (d) novel regulators of interferon responses (Fig. 3F). From this analysis, we identified USP2 as a candidate activator DUB (DUB^A^) and OTUD5 as a candidate inhibitor DUB (DUB^I^) of IRF7 transcriptional activity (Fig. 3F, boxed), and selected them for further mechanistic studies.

### USP2 Functions as a Positive Regulator and OTUD5 as a Negative Regulator of IRF7 Transcriptional Activity

To elucidate the mechanisms by which USP2 and OTUD5 regulate IRF7 transcriptional activity, we employed RAW264.7 macrophages—a murine myeloid cell line that expresses endogenous IRF7. Upon stimulation with SeV or poly(I:C) (a synthetic analog of viral RNA that activates RLR and TLR3 pathways), RAW264.7 cells showed robust upregulation of *Otud5* mRNA (Fig. 4A, B). To assess the functional relevance of OTUD5 and USP2 in these cells, we performed siRNA-mediated knockdown (Fig. 4B, C). As predicted from our screen, knockdown of *Otud5* significantly enhanced the expression of IRF7-dependent antiviral genes, including *Ifna*, *Ifnb1*, *Ifit1*, and *Ifit3*, compared to non-targeting (NT) control siRNA (Fig. 4D–G). Conversely, knockdown of *Usp2* led to a marked reduction in the expression of the same set of genes (Fig. 4D–G), supporting the conclusion that USP2 functions as a positive regulator (DUB^A^) and OTUD5 as a negative regulator (DUB^I^) of IRF7-mediated transcription. To confirm that these effects were independent of IRF3, which is also expressed in RAW264.7 cells, we validated the findings in the HEK-KO.IRF7 cell model that lacks IRF3. In these cells, knockdown of USP2 suppressed, while knockdown of OTUD5 enhanced, *IFNA* expression following viral stimulation (Fig. 4H–J), consistent with the observations in macrophages. Together, these results confirm that USP2 promotes, and OTUD5 suppresses, IRF7 transcriptional activity, validating their roles as DUB^A^ and DUB^I^, respectively, in both human and murine cellular contexts.

**Figure 4.**
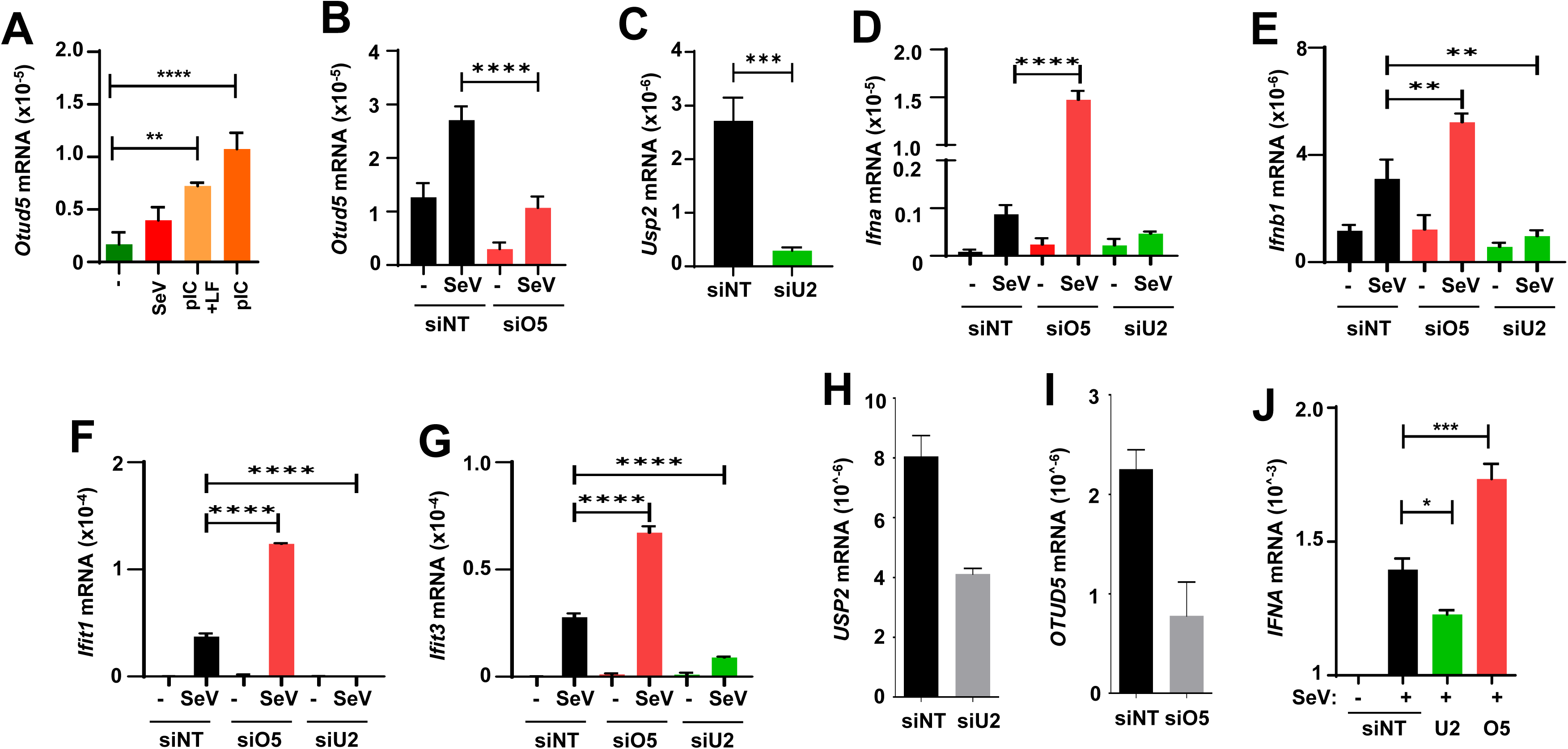
USP2 and OTUD5 differentially regulate IRF7 activity in myeloid and non-myeloid cells. **(A)** RAW-Luci macrophages were either infected with SeV, or transfected with poly(I:C) (pIC) using Lipofectamine (LF), or treated with pIC alone. *Otud5* mRNA levels were quantified by qRT-PCR. **(B–G)** RAW264.7 cells were transfected with NT, Otud5 (O5), or Usp2 (U2) siRNAs, infected with SeV for 8 hours, and mRNA levels of *Otud5, Usp2, Ifna, Ifnb1, Ifit1,* and *Ifit3* were analyzed by qRT-PCR. **(H–J)** HEK-KO.IRF7 cells were transfected with NT, OTUD5 (O5), or USP2 (U2) siRNAs and infected with SeV. *USP2*, *OTUD5*, and *IFNA* mRNA levels were measured by qRT-PCR. The data represent mean ± SEM (A-G, J), * p<0.05, ** p<0.001, *** p<0.001, **** p<0.0001.

### USP2 and OTUD5 Interact with IRF7 During Viral Infection

Given their roles as regulators of IRF7 transcriptional activity, we investigated whether USP2 and OTUD5 physically interact with IRF7 during antiviral responses. Using HEK-KO.IRF7 cells, we performed co-immunoprecipitation (co-IP) to assess the interaction between endogenous USP2 and IRF7. USP2 showed a low basal association with IRF7 in uninfected cells, which markedly increased upon SeV infection (Fig. 5A). To validate this interaction in primary immune cells, we performed proximity ligation assays (PLA) in bone marrow-derived macrophages (BMDMs). SeV infection resulted in a robust increase in PLA signal for USP2– IRF7 interaction, confirming infection-induced complex formation in a physiologically relevant context (Fig. 5B, C). We next examined the interaction between IRF7 and OTUD5. Similar to USP2, OTUD5 co-immunoprecipitated with IRF7 in HEK-KO.IRF7 cells during SeV infection (Fig. 5D). PLA analysis in SeV-infected BMDMs further confirmed a strong increase in IRF7–OTUD5 interaction (Fig. 5E, F). These findings indicate that both USP2 and OTUD5 interact with IRF7 in response to viral infection in myeloid and non-myeloid cells.

**Figure 5.**
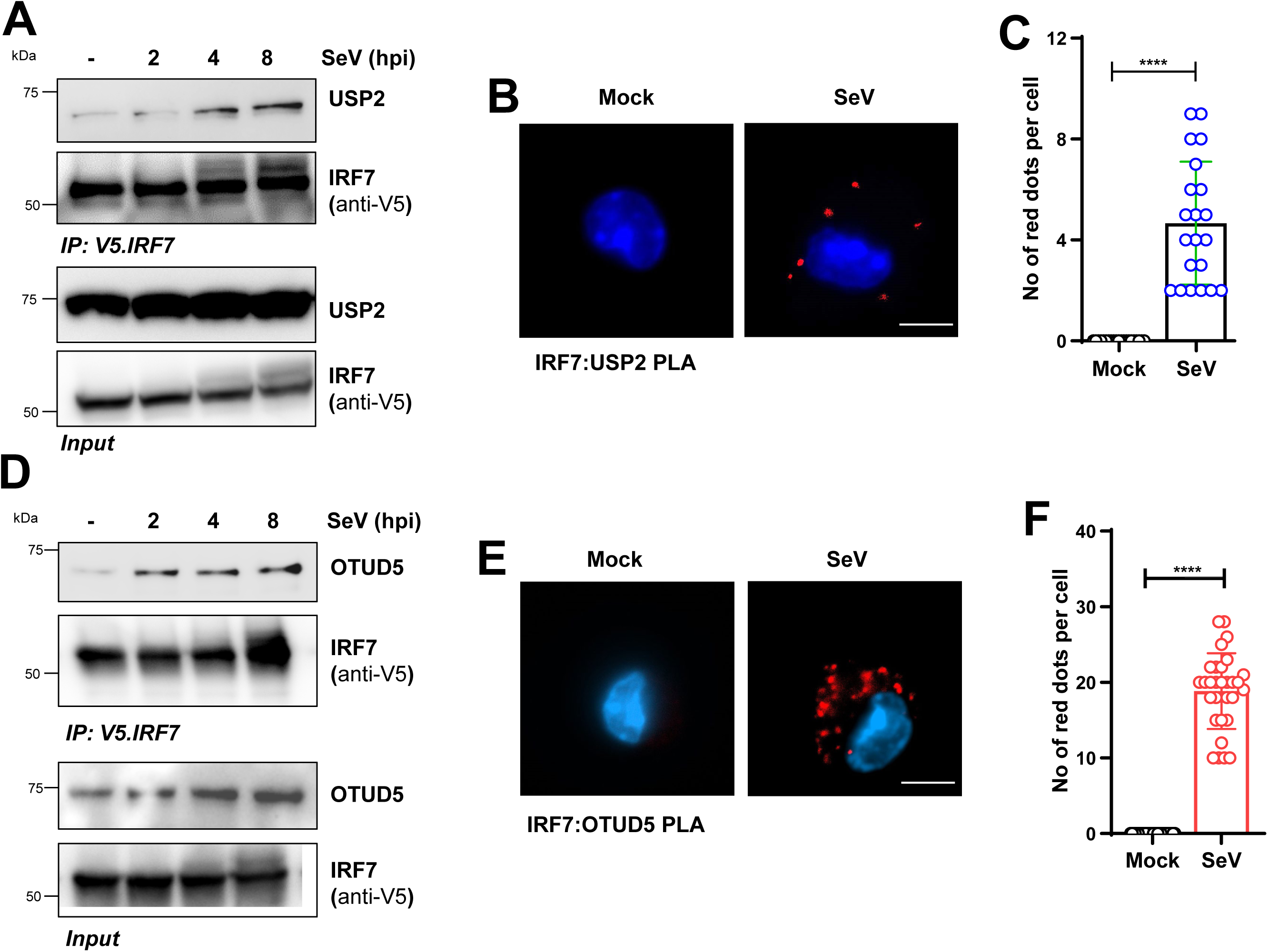
IRF7 interacts with USP2 and OTUD5 upon SeV infection. **(A)** USP2:IRF7 interaction in HEK-KO.IRF7 cells was assessed by co-immunoprecipitation at the indicated times post-SeV infection. **(B–C)** Proximity ligation assay (PLA) was performed in WT primary BMDMs using anti-USP2 and anti-IRF7 antibodies in mock or SeV-infected (8 hpi) cells. Red dots represent interaction signals, quantified in (C). **(D)** OTUD5:IRF7 interaction in HEK-KO.IRF7 cells was analyzed by co-immunoprecipitation following SeV infection. **(E–F)** PLA was conducted in WT BMDMs using anti-OTUD5 and anti-IRF7 antibodies in mock or SeV-infected cells. Interaction signals were quantified in (F).

### USP2 and OTUD5 Regulate IRF7 Through Linkage-Specific Deubiquitination

To explore whether these interactions affect IRF7 ubiquitination, we examined the specificity of USP2 and OTUD5 for different ubiquitin linkages on IRF7 by ectopically expressing them in HEK-KO.IRF7 cells (Fig. 6A). Since OTUD5 acts as a negative regulator of IRF7 activity, we first tested its effect on K63-linked ubiquitination, a modification known to activate IRF7. In HEK-KO.IRF7 cells expressing the K63-only ubiquitin mutant (Ub-K63O), OTUD5 overexpression led to strong suppression of K63-linked ubiquitinated IRF7 (Ub^63^-IRF7), in both uninfected and SeV-infected conditions (Fig. 6B). Next, we tested the effects of USP2 and OTUD5 on K27- and K33-linked ubiquitination using Ub-K27O and Ub-K33O mutants. USP2, but not OTUD5, selectively reduced K27-linked ubiquitination of IRF7 (Ub^27^-IRF7), whereas neither enzyme affected Ub^33^-IRF7 (Fig. 6C, D). These results establish the linkage specificity of these DUBs: USP2 targets Ub^27^-IRF7 and OTUD5 targets Ub^63^-IRF7. To map the sites of these modifications, we used IRF7 truncation mutants and performed ubiquitination assays. Both Ub^27^- and Ub^63^-linked IRF7 levels were significantly reduced in the IRF7(1–237) mutant compared to the IRF7(1–410) construct (Fig. 6E-G), implicating the C-terminal region in ubiquitin modification. Based on our results from the IRF7 deletion mutants and prior knowledge of Ub^63^ sites, we postulated that lysines K327 and K329 are candidate Ub^27^ sites (Fig. 6H). We then generated IRF7 lysine-to-arginine mutants—K327R (K1), K329R (K2), and the double mutant KK327/329RR (K3)—to assess their contributions to Ub^27^-IRF7 linkage. SeV infection induced Ub^27^-IRF7 in HEK-KO.IRF7 cells expressing wild-type IRF7, while all three KR mutants exhibited significantly reduced Ub^27^-IRF7 levels (Fig. 6I, J). These results identify K327 and K329 as the principal sites for K27-linked ubiquitination of IRF7 during viral infection and establish USP2 and OTUD5 as selective regulators of Ub^27^- and Ub^63^-linked IRF7, respectively.

**Figure 6.**
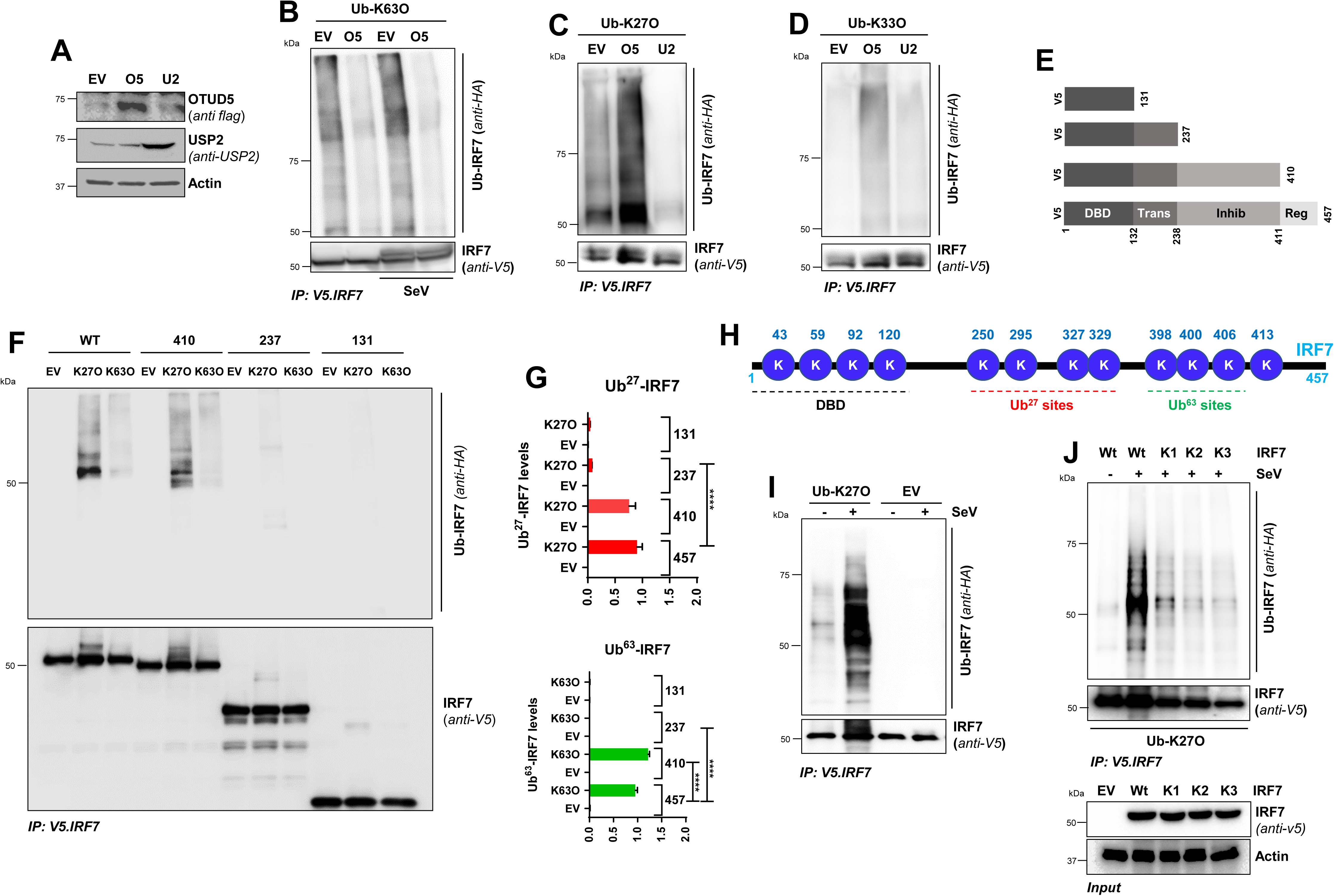
OTUD5 and USP2 regulate distinct ubiquitin linkages on IRF7. **(A)** HEK-KO.IRF7 cells were transfected with Flag.OTUD5 or USP2 plasmids, and the cell lysates were analyzed by immunoblot, as indicated. **(B–D)** HEK-KO.IRF7 cells were transfected with HA-tagged Ub-K63O, Ub-K27O, or Ub-K33O plasmids, in the absence or the presence of OTUD5 or USP2, as indicated, and infected with SeV. The cell lysates were immunoprecipitated with anti-V5 antibody and immunoblotted with anti-HA, as indicated. **(E–G)** Full-length and truncated IRF7 mutants (E) were co-transfected with Ub-K63O or Ub-K27O, followed by SeV infection. Ub-IRF7 levels were measured by IP and immunoblot (F) and quantified using ImageJ (G). **(H)** Schematic of IRF7 protein domains and putative ubiquitin linkage sites (K63 and K27); DBD, DNA-binding domain. **(I)** Cells were transfected with HA.Ub-K27O and infected with SeV. Ub-IRF7 was analyzed as in (A). **(J)** IRF7-WT or KR mutants (K1: K327R, K2: K329R, K3: K327/329RR) were co-transfected with Ub-K27O, infected with SeV, and analyzed for Ub-IRF7. The lower panel shows the expression of the IRF7 mutants. EV, empty vector; Ub-IRF7, ubiquitinated IRF7.

### K27-Linked Ubiquitination Inhibits IRF7 Activation and Transcriptional Function

Given the novel Ub^27^-mediated regulation of IRF7 activity, we focused on dissecting the molecular mechanism. To investigate the functional impact of K27-linked ubiquitination on IRF7 activation, we analyzed the levels of phosphorylated IRF7 (p-IRF7) in HEK-KO.IRF7 cells expressing either Ub-K27O or Ub-K63O. SDS-PAGE analysis revealed that Ub-K63O expression enhanced the lower mobility p-IRF7 band, while Ub-K27O suppressed it (Fig. 7A, B). We confirmed these findings by examining pSer425/426, key phosphorylation sites required for IRF7 activation, in nuclear extracts. Expression of Ub-K27O led to decreased pSer-IRF7 and accumulation of unphosphorylated IRF7 in the nucleus (Fig. 7C, D), suggesting that K27-linked ubiquitination inhibits IRF7 phosphorylation and nuclear activation. To directly assess transcriptional activity, we analyzed IRF7 target gene expression. Enrichment of Ub-K27O suppressed the induction of *IFNB1*, *IFIT1*, *IFIT2*, and *IFIT3* mRNAs (Fig. 7E–H), indicating that K27-linked ubiquitination impairs IRF7-driven antiviral gene expression. Finally, we evaluated the transcriptional activity of the K1 and K2 IRF7 mutants, which are defective in Ub^27^ modification. Using the HEK-SEAP-KO reporter cell line expressing an ISRE-driven SEAP construct, both mutants exhibited significantly higher SEAP activity than wild-type IRF7 (Fig. 7I). Notably, Ub-K27O suppressed SEAP activity in wild-type IRF7-expressing cells but had no effect on K1 or K2 mutants (Fig. 7I), further demonstrating that K27-linked ubiquitination at K327 and K329 suppresses IRF7 transcriptional activity.

**Figure 7.**
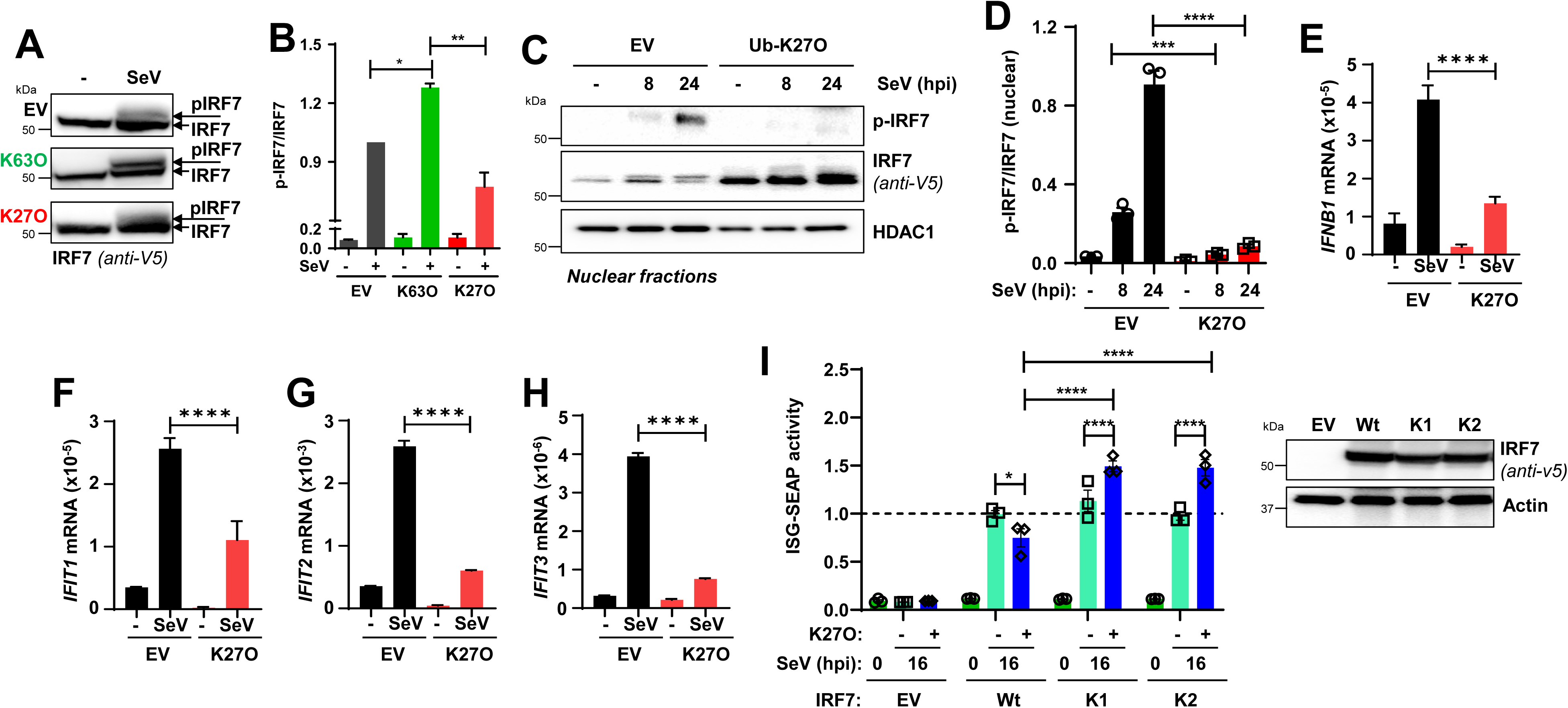
K27-linked ubiquitination suppresses IRF7 phosphorylation and transcriptional activity. (A–B) HEK-KO.IRF7 cells were transfected with Ub-K63O or Ub-K27O plasmids, infected with SeV, and IRF7 phosphorylation (p-IRF7) was analyzed by immunoblot (A). Quantification of p-IRF7 and total IRF7 was performed using Image J (B). **(C–D)** Nuclear fractions from cells transfected with Ub-K27O and infected with SeV were analyzed for p-IRF7, total IRF7, and HDAC1. Nuclear p-IRF7 was quantified in (D). **(E–H)** IRF7-WT and Ub-K27O were co-transfected in HEK-KO cells, followed by SeV infection. Expression of *IFNB1*, *IFIT1*, *IFIT2*, and *IFIT3* was measured by qRT-PCR. **(I)** IRF3-deficient HEK293.ISG-SEAP (HEK.SEAP-KO) cells were co-transfected with IRF7 (WT, K1, or K2) and Ub-K27O, infected with SeV, and SEAP activity was measured. IRF7 protein expression is shown in the right panel. EV, empty vector.

## Discussion

IRF7, a key member of the IRF family and the so-called “master transcription factor” of type I interferons (IFNs), is predominantly expressed in myeloid and lung epithelial cells (30, 31). Its expression is further induced by viral infection or tissue damage. In resting cells, IRF7 resides in the cytosol in an inactive state. Activation of pattern recognition receptors (PRRs) triggers IRF7 transcriptional activity, which is tightly regulated by post-translational modifications (8, 31). Phosphorylation of conserved serine residues (Ser425/426 or Ser437/438), primarily mediated by TBK1 or IKKε, leads to IRF7 nuclear translocation and gene induction. In addition to phosphorylation, ubiquitination plays a pivotal role in modulating IRF7 activity. K63-linked ubiquitination of IRF7, catalyzed by E3 ligases such as TRAF6 and NEURL3, promotes its transcriptional function (15, 32). However, ubiquitination is reversible and subject to regulation by deubiquitinases (DUBs), which remove specific ubiquitin chains to maintain cellular homeostasis. The interplay between phosphorylated IRF7 (p-IRF7) and ubiquitinated IRF7 (Ub-IRF7) remains incompletely understood.

To dissect this regulatory axis, we conducted an unbiased genetic screen to identify IRF7-specific DUBs. Our screen uncovered two DUBs with opposing roles: OTUD5 and USP2. OTUD5 functioned as a negative regulator, interacting with IRF7, removing K63-linked ubiquitin chains, and suppressing IRF7-dependent gene expression. In contrast, USP2 enhanced IRF7 activity by deubiquitinating K27-linked IRF7, thereby facilitating transcriptional activation. Importantly, we found that K27-linked ubiquitination inhibits IRF7 phosphorylation, suggesting that K27-Ub acts as a negative regulatory linkage. Together, these findings suggest a model in which IRF7 recruits both inhibitory and activating DUBs to fine-tune its activity.

Mechanistically, our data suggest that IRF7 exists in a K27-linked ubiquitinated form under homeostatic conditions, maintaining the transcription factor in an inactive state. Upon stimulation, USP2 removes K27-linked chains, potentially allowing TRAF6 to catalyze activating K63-linked ubiquitination, thereby promoting IRF7 activity. Subsequently, OTUD5 may be recruited to remove K63-linked chains and restore IRF7 to an inactive state. Notably, OTUD5 recruitment occurs early during infection, possibly acting through an indirect mechanism to reinforce K27-linkage. The identity of the E3 ligase responsible for K27-linked ubiquitination of IRF7 remains unknown. However, studies in zebrafish suggest that FBXO3 may catalyze this modification; FBXO3-deficient zebrafish are resistant to viral infection, supporting its role as a negative regulator of IRF7 (33). Whether FBXO3 serves a similar function in mammalian cells remains to be determined. Our findings also raise the possibility that mammalian IRF3 may undergo similar K27-linked regulation, given that FBXO3 modifies zebrafish IRF3 and that USP2 could serve as a shared regulatory DUB. These insights reveal a previously unappreciated noncanonical ubiquitin modification that negatively regulates IRF7 function. Further investigation is needed to determine whether K27-linked IRF7 plays roles in other physiological or pathological contexts, such as autoimmune diseases, where IRF7 is a critical contributor. Although K63-linked ubiquitination of IRF7 has been previously reported, most notably in Epstein-Barr Virus (EBV)-infected cells where LMP1 recruits TRAF6 to form a positive feedback loop involving IRF7 activation—our study demonstrates that this modification is subject to negative regulation by OTUD5 (15). Similarly, NEURL has recently been identified as a K63-specific E3 ligase promoting IRF7 antiviral function; NEURL-deficient mice exhibit increased viral susceptibility, underscoring the importance of K63-linked IRF7 in antiviral defense (32).

DUBs are emerging as central regulators of immune signaling. Their dysregulation can lead to immune dysfunction, inflammatory disease, and cancer (30, 34). However, how specific DUBs are activated and recruited to IRF7 remains unclear. Prior studies, primarily in zebrafish, have implicated other DUBs such as USP8, which promotes autophagy-dependent degradation of IRF7 during SVCV infection, and A20, which suppresses IRF7 activity in EBV-infected cells (19, 35). To identify IRF7-specific regulators in human cells, we performed a siRNA-based screen in a system lacking IRF3, thereby isolating IRF7-driven responses. Although we characterized OTUD5 and USP2 in depth, other candidate DUBs from our screen (see Fig. 3F) may also regulate IRF7 and warrant future investigation. Our data indicate that OTUD5 expression is upregulated by viral infection or PRR stimulation, suggesting it acts as a feedback inhibitor in antiviral signaling. We also observed increased IRF7–OTUD5 binding following SeV infection. In contrast, USP2 regulation is poorly understood; future studies are needed to clarify how its expression or activity is controlled. Several critical questions remain. What determines the specificity of IRF7–DUB interactions? In which cellular compartments do these interactions occur? How are DUBs temporally regulated during infection? Addressing these gaps will deepen our understanding of how IRF7 activity is dynamically regulated through ubiquitination.

In summary, our study reveals that IRF7 activity is regulated by distinct ubiquitin linkages— K63-linked ubiquitination promotes transcriptional activation, while K27-linked ubiquitination inhibits it. Two DUBs, OTUD5 and USP2, selectively remove these linkages to suppress or enhance IRF7 activity, respectively. These findings reveal a new layer of IRF7 regulation and lay the groundwork for exploring DUBs as therapeutic targets in viral and autoimmune diseases where IRF7 plays a central role.

